# The role of group I p21-activated kinases in contraction-stimulated skeletal muscle glucose transport

**DOI:** 10.1101/2020.01.29.925024

**Authors:** Lisbeth L. V. Møller, Ida L. Nielsen, Jonas R. Knudsen, Nicoline R. Andersen, Thomas E. Jensen, Lykke Sylow, Erik A. Richter

## Abstract

**Aim:** Muscle contraction stimulates skeletal muscle glucose transport. Since it occurs independently of insulin, it is an important alternative pathway to increase glucose uptake in insulin-resistant states, but the intracellular signalling mechanisms are not fully understood. Muscle contraction activates group I p21-activated kinases (PAKs) in mouse and human skeletal muscle. PAK1 and PAK2 are downstream targets of Rac1, which is a key regulator of contraction-stimulated glucose transport. Thus, PAK1 and PAK2 could be downstream effectors of Rac1 in contraction-stimulated glucose transport. The current study aimed to test the hypothesis that PAK1 and/or PAK2 regulate contraction-induced glucose transport.

**Methods:** Glucose transport was measured in isolated soleus and extensor digitorum longus (EDL) mouse skeletal muscle incubated either in the presence or absence of a pharmacological inhibitor (IPA-3) of group I PAKs or originating from whole-body PAK1 knockout (KO), muscle-specific PAK2 (m)KO or double whole-body PAK1 and muscle-specific PAK2 knockout mice.

**Results:** IPA-3 attenuated (−22%) the increase in muscle glucose transport in response to electrically-stimulated contraction. PAK1 was dispensable for contraction-stimulated glucose uptake in both soleus and EDL muscle. Lack of PAK2, either alone (−13%) or in combination with PAK1 (−14%), reduced contraction-stimulated glucose transport compared to control littermates in EDL, but not soleus muscle.

**Conclusion:** Contraction-stimulated glucose transport in isolated glycolytic mouse EDL muscle is partly dependent on PAK2, but not PAK1.

## Introduction

Muscle contraction increases skeletal muscle glucose uptake independently of insulin ^1–3^. Accordingly, muscle contraction increases glucose uptake in both insulin-sensitive and insulin-resistant skeletal muscle ^4–6^. Additionally, insulin sensitivity is improved after cessation of muscle contraction ^7–10^, making muscle contraction during acute exercise a non-pharmacological treatment for insulin resistance ^11^. However, while muscle contraction is known to promote the translocation of the glucose transporter (GLUT)-4 to the plasma membrane, which facilitates glucose entry into the muscle, the intracellular signalling regulating this process is not completely understood.

Upon muscle contraction, multiple intracellular signalling pathways are activated that promote GLUT4 translocation and a subsequent increase in muscle glucose transport. Redundant Ca^+2^-dependent signalling, metabolic stress signalling, and mechanical stress signalling are proposed to regulate distinct steps important for glucose transport in response to muscle contraction ^12^. The group I p21-activated kinase (PAK)-1 and PAK2 are activated in response to electrical pulse stimulation in C2C12 myotubes ^13,14^ and muscle contraction/acute exercise in mouse and human skeletal muscle ^15^. Group I PAKs (PAK1-3) are downstream targets of the Rho family GTPases Cdc42 and Rac1 ^16^. Rac1 plays a key role in mediating glucose uptake in response to muscle contraction and acute exercise in skeletal muscle ^15,17,18^. Additionally, the contraction-stimulated increase in PAK1/2 activity is blunted in muscles from muscle-specific Rac1 knock-out (KO) mice ^15^, suggesting a potential role for PAK1 and/or PAK2 in regulating muscle glucose uptake during muscle contraction. However, the significance of the increased activity of group I PAKs downstream of Rac1 in response to muscle contraction is unknown. We hypothesized that PAK1 and PAK2 participate in the regulation of glucose uptake in response to contraction, due to their well-described role as Rac1 effector proteins. Our results identify PAK2, but not PAK1, as a partial requirement for contraction-stimulated glucose transport in mouse skeletal muscle.

## Results

### Contraction-stimulated glucose transport is partially inhibited by pharmacological inhibition of PAK1/2

To investigate the role of group I PAKs in the regulation of contraction-stimulated glucose transport, we first analyzed 2-deoxyglucose (2DG) transport in isolated soleus and extensor digitorum (EDL) muscle in the presence or absence of a pharmacological group I PAK inhibitor, IPA-3. Contractions increased 2DG transport in DMSO-treated soleus (2.9-fold) and EDL (3.0-fold) muscle (Fig. 1A+B). IPA-3 partly inhibited contraction-stimulated 2DG transport in soleus (−22%) and EDL (−22%; Fig. 1A+B). The reduction in contraction-stimulated 2DG transport upon IPA-3 treatment was not associated with reduced initial force development in IPA-3 treated muscles (Fig 1C). While phosphorylated (p)AMPKα T172 was unaffected by IPA-3 in soleus muscle (Fig. 1D), contraction-stimulated pAMPKα T172 was reduced (−46%) in IPA-3-treated EDL muscle (Fig. 1E). However, AMPKs downstream target pACC1/2 S79/212 was normally phosphorylated in response to contraction in both muscles (Fig. 1F+G), suggesting that the AMPK-ACC signalling pathway was largely unaffected by IPA-3 treatment. Altogether, these data suggest that contraction-stimulated glucose transport partly relies on group I PAKs in skeletal muscles.

**Figure 1:**
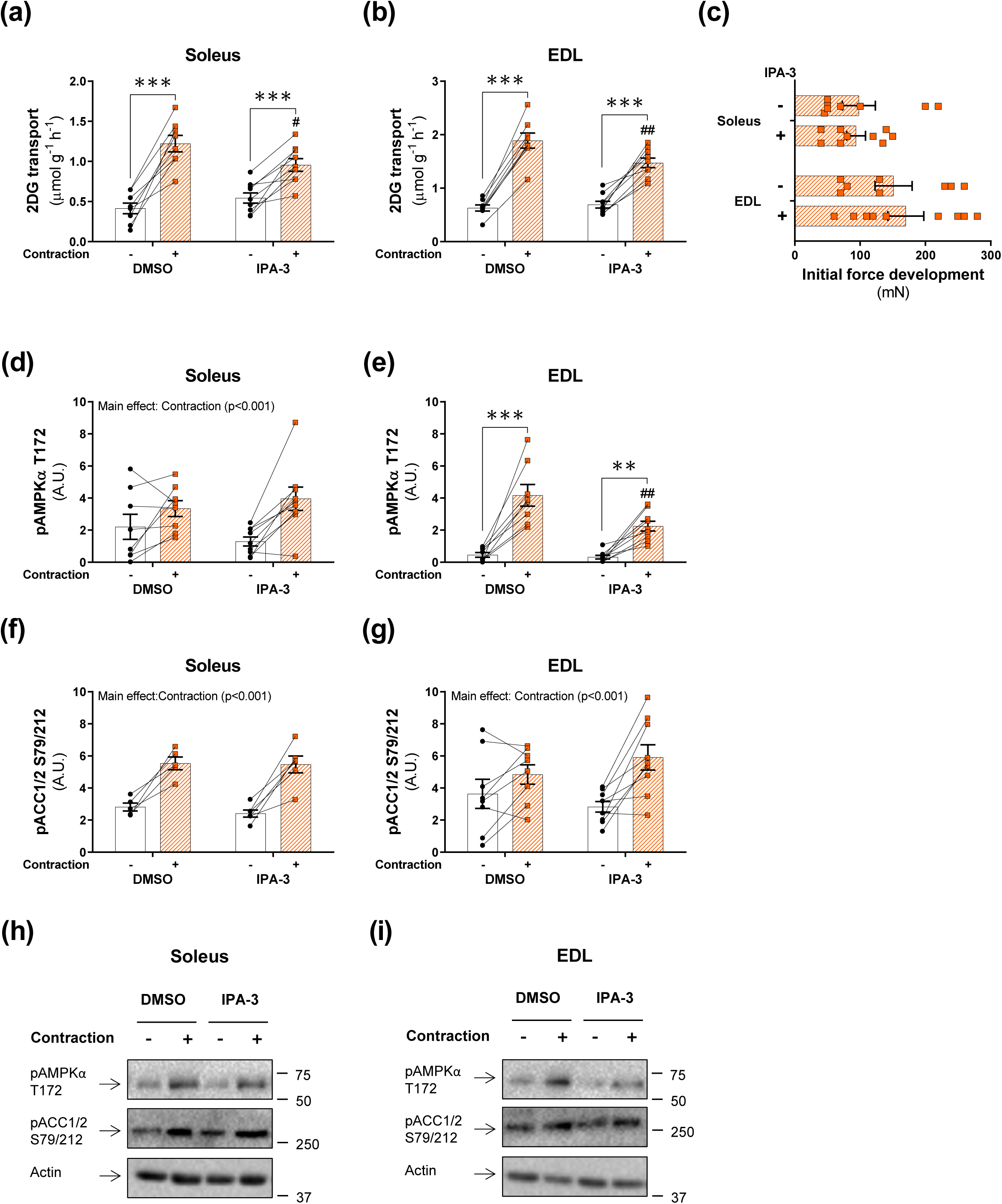
Contraction-stimulated glucose transport is partially inhibited by pharmacological inhibition of PAK1/2. **(a-b)** Contraction-stimulated (2 sec/15 sec, 100 Hz) 2-deoxyglucose (2DG) transport in isolated soleus (a) and extensor digitorum longus (EDL; b) muscle ± 40 µM IPA-3 or a corresponding amount of DMSO (0.11%). Isolated muscles were pre-incubated for 45 minutes followed by 15 minutes of electrically-stimulated contraction with 2DG transport measured for the final 10 minutes of stimulation. Data were evaluated with a two-way repeated measures (RM) ANOVA. **(c)** Initial force development during electrically-stimulated contraction. Data were evaluated with a Student’s t-test. **(d-g)** Quantification of phosphorylated (p)AMPKα T172 and pACC1/2 S79/212 in contraction-stimulated soleus (d and f) and EDL (e and g) muscle. Data were evaluated with a two-way RM ANOVA. Some of the data points were excluded due to the quality of the immunoblot, and the number of determinations was *n = 5/6* (DMSO/IPA-3) for pACC1/2 S79/212 in soleus muscle. **(h-i)** Representative blots showing pAMPKα T172 and pACC S212 and actin protein expression as a loading control in soleus (h) and EDL (i) muscle. Main effects are indicated in the panels. Interactions in two-way RM ANOVA were evaluated by Tukey’s post hoc test: Contraction vs. basal **/*** (p<0.01/0.001); IPA-3 vs. DMSO ## (p<0.01). Unless stated previously in the figure legend, the number of determinations in each group: Soleus, *n = 8/9* (DMSO/IPA-3); EDL, *n = 8/9*. Data are presented as mean ± S.E.M. with individual data points shown. Paired data points are connected with a straight line. A.U., arbitrary units.

### Contraction-stimulated glucose transport partially relies on PAK2, but not PAK1, in mouse EDL muscle

IPA-3 is a pharmacological inhibitor of group I PAKs (PAK1-3) of which PAK1 and PAK2 are detectable in skeletal muscle ^19–21^. To identify the relative role of PAK1 and PAK2 in the regulation of contraction-stimulated glucose transport, we investigated contraction-stimulated glucose transport in isolated soleus and EDL muscles from a cohort of PAK1 KO, PAK2 mKO, and double knockout mice with whole-body knockout of PAK1 and muscle-specific knockout of PAK2 (1/m2 dKO) compared to control littermates (Fig. 2A+B). The whole-body metabolic characteristics of this cohort of mice have previously been described ^22^.

**Figure 2:**
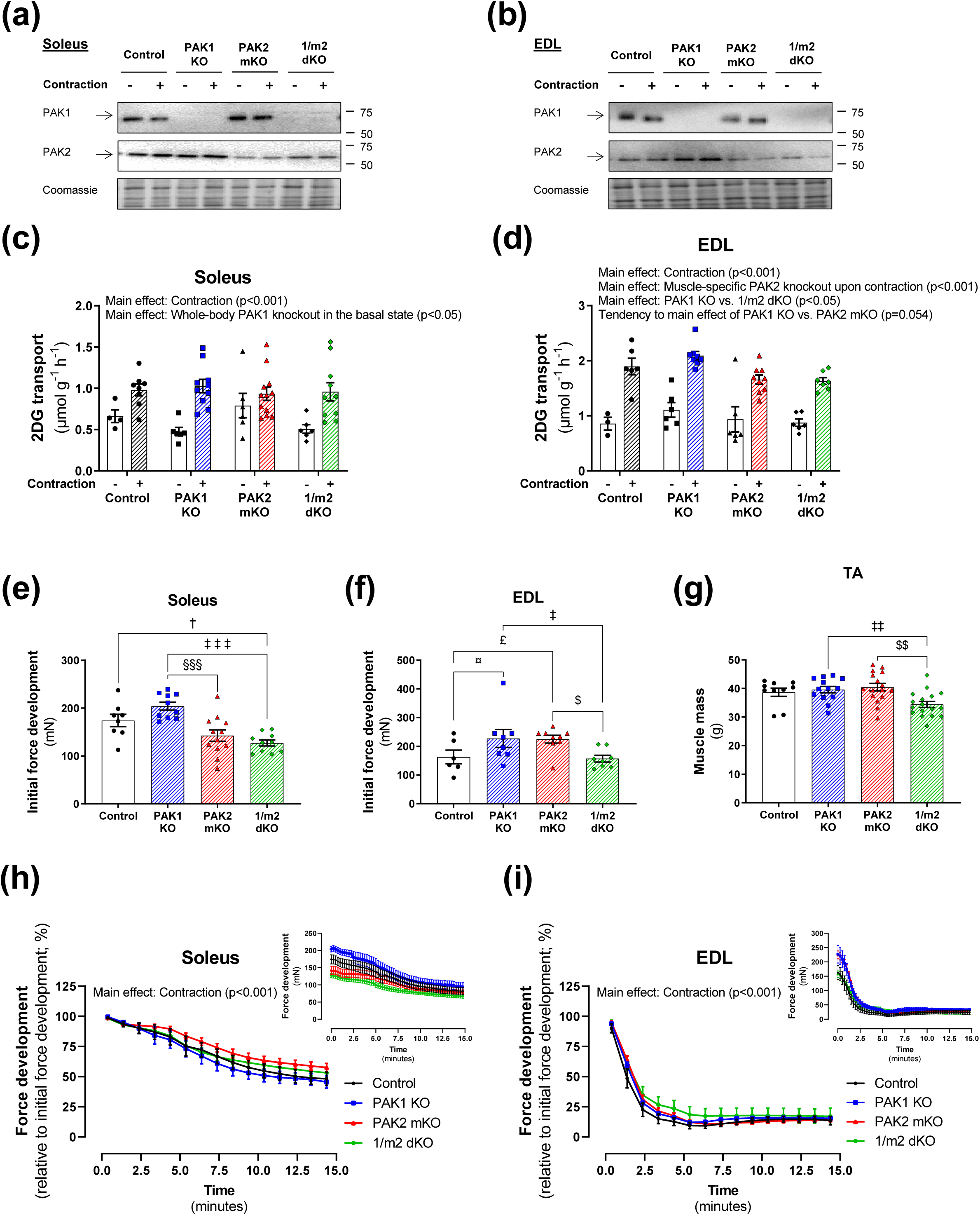
Contraction-stimulated glucose transport partially requires PAK2, but not PAK1, in mouse EDL muscle. **(a-b)** Representative blots showing PAK1 and PAK2 protein expression in soleus (a) and extensor digitorum longus (EDL; b) muscle from whole-body PAK1 knockout (KO), muscle-specific PAK2 (m)KO, PAK1/2 double KO (1/m2 dKO) mice or control littermates. **(c-d)** Contraction-stimulated (2 sec/15 sec, 100 Hz) 2-deoxyglucose (2DG) transport in isolated soleus (c) and EDL (d) muscle from PAK1 KO, PAK2 mKO, 1/m2 dKO mice or control littermates. Isolated muscles were pre-incubated for 10-20 minutes followed by 15 minutes of electrically-stimulated contraction with 2DG transport measured for the final 10 minutes of stimulation. The number of determinations in each group for soleus: Control, *n = 4/8* (Basal/Contraction); PAK1 KO, *n = 6/10*; PAK2 mKO, *n = 6/12*; 1/m2 dKO, *n = 6/10*, and for EDL: Control, *n = 3/6* (Basal/Contraction); PAK1 KO, *n = 6/8*; PAK2 mKO, *n = 6/9*; 1/m2 dKO, *n = 6/7*. Data were evaluated with two twoway ANOVAs to test the factors ‘PAK1’ (PAK1^+/-^ vs. PAK^-/-^) and ‘PAK2’ (PAK2^fl/fl^;MyoD^+/+^ vs. PAK2^fl/fl^;MyoD^iCre/+^) in the basal and contraction-stimulated state, respectively, thereby assessing the relative contribution of PAK1 and PAK2. Differences between genotypes and the effect of contraction were assessed with a two-way ANOVA to test the factors ‘Genotype’ (Control vs. PAK1 KO vs. PAK2 mKO vs. d1/2 KO) and ‘Stimuli’ (Basal vs. Contraction). **(e-f)** Initial force development during electrically-stimulated contractions in soleus (e) and EDL (f) muscle. The number of determinations in each group: Control, *n = 8/6* (soleus/EDL); PAK1 KO, *n = 10/8*; PAK2 KO, *n = 12/9*; 1/m2 dKO, *n = 10/8*. Data were evaluated with a two-way ANOVA to test the factors ‘PAK1’ and ‘PAK2’, thereby assessing the relative contribution of PAK1 and PAK2. Differences between genotypes were evaluated with a one-way ANOVA. **(g)** Tibialis anterior (TA) muscle mass in PAK1 KO, PAK2 mKO, 1/m2 dKO mice or control littermates. The number of determinations in each group: Control, *n = 10*; PAK1 KO, *n = 13*; PAK2 KO, *n = 16*; 1/m2 dKO, *n = 17*. Data were evaluated with a two-way ANOVA to test the factors ‘PAK1’ and ‘PAK2’, thereby assessing the relative contribution of PAK1 and PAK2. Differences between genotypes were evaluated with a one-way ANOVA. **(h-i)** Force development relative to initial force development in soleus (H) and EDL (I) muscle from whole-body PAK1 KO, PAK2 mKO, 1/m2 dKO mice or control littermates. Data points relative to initial force development is an average of the values at four consecutive time points. Original force development data inserted in the upper right corner. The number of determinations in each group: Control, *n = 8/6* (soleus/EDL); PAK1 KO, *n = 10/8*; PAK2 KO, *n = 12/9*; 1/m2 dKO, *n = 10/8*. Due to missing data points, differences between genotypes and the effect of electrically-stimulated contraction were assessed with a mixed-effects model analysis to test the factors ‘Genotype’ and ‘Time point’. Main effects are indicated in the panels. Significant one-way ANOVA and interactions in two-way (RM when applicable) ANOVA were evaluated by Tukey’s post hoc test: Control vs. PAK1 KO ¤ (p<0.05); Control vs. PAK2 mKO £ (p<0.05); Control vs. d1/2 KO † (p<0.05); PAK1 KO vs. PAK2 mKO §§§ (p<0.001); PAK1 KO vs. d1/2 KO ‡/‡‡/‡‡‡ (p<0.05/0.01/0.001); PAK2 mKO vs. d1/2 KO $/$$ (p<0.05/0.01). Data are presented as mean ± S.E.M. with individual data points shown.

In soleus muscle, contraction-stimulated glucose transport was unaffected by the lack of PAK1, PAK2 or both PAKs combined (Fig. 2C). In contrast, in EDL lack of PAK2, either alone or in combination with PAK1 KO, partially reduced contraction-stimulated glucose transport compared to PAK1 KO mice (PAK2 mKO: −21%; 1/m2 dKO: −22%) and control littermates (PAK2 mKO: −13%; 1/m2 dKO: −14%; Fig. 2D). Lack of PAK2 decreased initial force development in soleus compared to PAK1 KO mice (PAK2 mKO: −30%; 1/m2 dKO: −38%) and control littermates (1/m2 dKO: −27%; Fig. 2E). In EDL, lack of PAK1 (+40%) or PAK2 (+38%) increased initial force development, while combined knockout of PAK1 and PAK2 decreased initial force development compared to PAK1 KO mice (−31%) and PAK2 mKO mice (−30%; Fig. 2F). The reduction in initial force development in 1/m2 dKO muscle could be ascribed to muscle wasting (−13%; Fig. 2G) as also previously reported for several distinct muscles in this mouse model ^20,23^. However, the decrease in force development over time was similar between all four genotypes in both soleus and EDL muscle (Fig. 2H+I). Thus, similar to insulin-stimulated glucose uptake ^22^, PAK1 is dispensable for contraction-stimulated glucose transport, while contraction-stimulated glucose transport partially relies on PAK2 in glycolytic EDL muscle.

### Canonical contraction signalling is largely unaffected by the lack of PAK1 and PAK2

Next, we investigated the effects of lack of PAK1 and/or PAK2 on contraction-stimulated molecular signalling. Lack of PAK2 tended (p=0.052) to reduce pAMPKα T172 in soleus (PAK2 mKO: −17%; 1/m2 dKO: −12%), but not EDL muscle (Fig. 3A+B). However, pACC1/2 S79/212 was normally phosphorylated in response to contractions in both muscles (Fig. 3C+D). Another contraction-stimulated downstream target of AMPKα2, pTBC1D1 S231 was unaffected by lack of PAK1 and/or PAK2 in soleus muscle (Fig. 3E), but was reduced (−39%) in 1/m2 dKO EDL muscle compared to muscle from PAK1 KO mice (Fig. 3F). Protein expression of AMPKα2, ACC and TBC1D1 was unaffected by lack of PAK1 and/or PAK2 (representative blots in Fig. 2K+L). We next analyzed the total protein content of proteins involved in glucose handling. Previously, in a slightly younger cohort (10-16 weeks of age vs. 26-35 weeks of age), we reported that GLUT4 protein expression was normal in soleus but mildly reduced in EDL in PAK2 mKO mice compared to littermate control ^22^. In contrast, GLUT4 protein expression was presently reduced in 1/m2 dKO soleus muscle from soleus compared to control muscle (−29%; Fig. 3G). In EDL muscle GLUT4 protein expression was unaffected by lack of PAK1 and/or PAK2 (Fig. 3H). Protein expression of hexokinase II (HKII), a key enzyme converting glucose to glucose-6-phosphate after uptake, was unaffected in soleus muscle (Fig. 3I), while higher (+34%) in 1/m2 dKO EDL muscle compared to PAK2 mKO muscle (Fig. 3J). Taken together, the reduced contraction-stimulated glucose transport in 1/m2 dKO EDL muscle was accompanied by impaired pTBC1D1 S237 phosphorylation (potentially decreasing glucose uptake) but also upregulation of HKII (potentially enhancing capacity for glucose uptake although a previous study suggest that in isolated muscles, HKII overexpression is not sufficient to increase neither basal nor insulin-stimulated glucose transport ^24^). Thus, the mechanism/s by which genetic ablation of PAK2 reduces contraction-stimulated glucose transport remain unclear.

**Figure 3:**
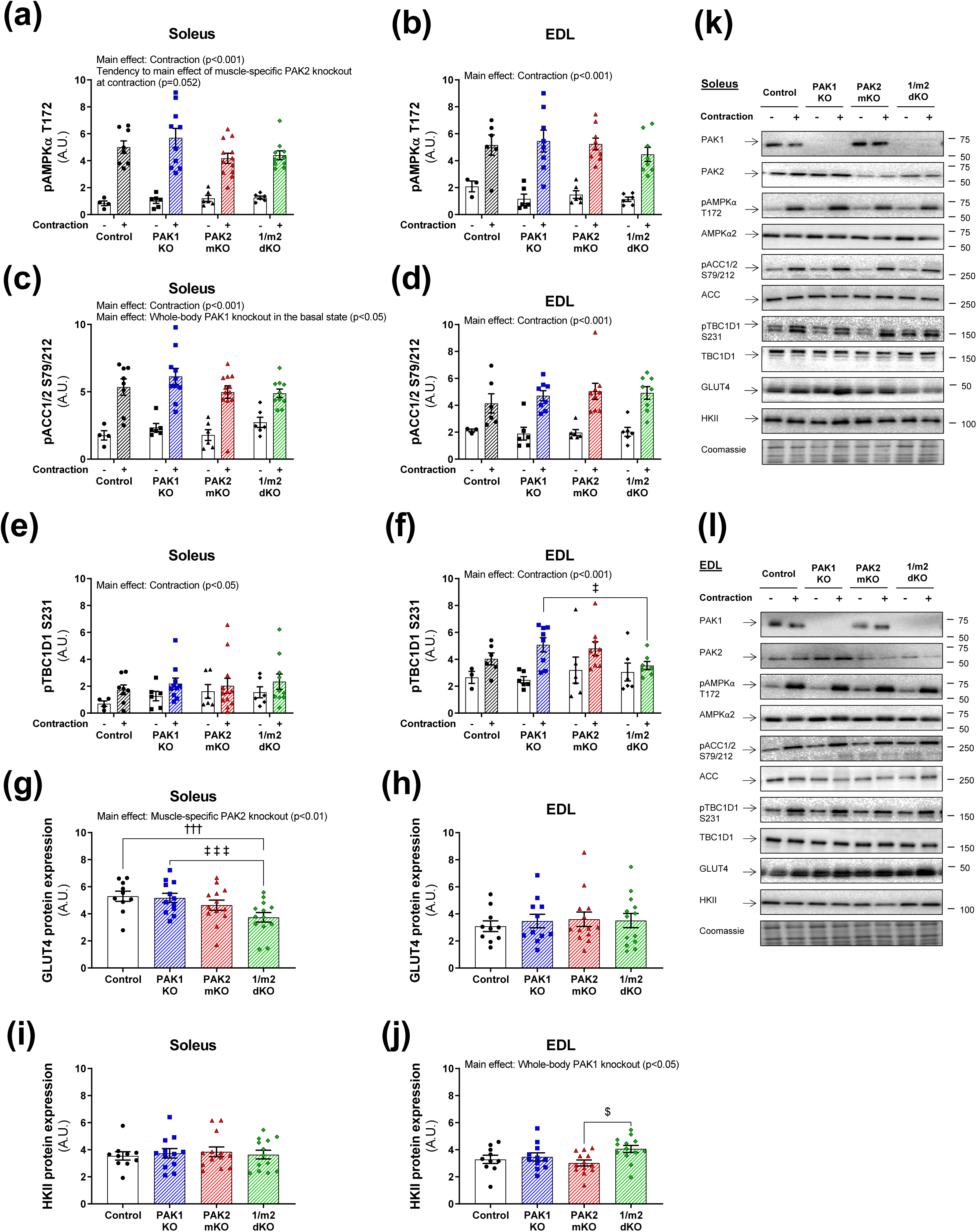
Canonical contraction signalling is largely unaffected by the lack of PAK1 and PAK2. **(a-j)** Quantification of phosphorylated (p)AMPKα T172, pACC1/2 S79/212, pTBC1D1 S231 and total GLUT4 and HKII protein expression in response to electrically-stimulated contractions (2 sec/15 sec, 100 Hz) in soleus (a, c, e, g, and i) and extensor digitorum longus (EDL; b, d, f, h, and j) muscle from whole-body PAK1 knockout (KO), muscle-specific PAK2 (m)KO, PAK1/2 double KO (1/m2 dKO) mice or control littermates. Total protein expression is an average of the two muscles from the same mouse. Protein phosphorylation was evaluated with two two-way ANOVAs to test the factors ‘PAK1’ (PAK1^+/-^ vs. PAK^-/-^) and ‘PAK2’ (PAK2^fl/fl^;MyoD^+/+^ vs. PAK2^fl/fl^;MyoD^iCre/+^) in basal and contraction-stimulated samples, respectively, thereby assessing the relative contribution of PAK1 and PAK2. Differences between genotypes and the effect of contraction stimulation were assessed with a two-way ANOVA to test the factors ‘Genotype’ (Control vs. PAK1 KO vs. PAK2 mKO vs. 1/m2 dKO) and ‘Stimuli’ (Basal vs. Contraction). Total protein expression was evaluated with a two-way ANOVA to test the factors ‘PAK1’ and ‘PAK2’ thereby assessing the relative contribution of PAK1 and PAK2, respectively. Differences between genotypes were evaluated with a one-way ANOVA. **(k-l)** Representative blots showing pAMPKα T172, pACC1/2 S79/212, pTBC1D1 S231 and total AMPKα2, ACC, TBC1D1, GLUT4 and HKII protein expression and coomassie staining as a loading control in soleus (k) and EDL (l) muscle. Main effects are indicated in the panels. Significant one-way ANOVA and interactions in two-way ANOVA were evaluated by Tukey’s post hoc test: Control vs. d1/2 KO ††† (p<0.001); PAK1 KO vs. d1/2 KO ‡/‡‡‡ (p<0.05/0.001); PAK2 mKO vs. d1/2 KO $ (p<0.05). The number of determinations in each group for soleus: Control, *n = 4/8* (Basal/Contraction); PAK1 KO, *n = 6/10*; PAK2 mKO, *n = 6/12*; 1/m2 dKO, *n = 6/10*. The number of determinations in each group for EDL: Control, *n = 3/6* (Basal/Contraction); PAK1 KO, *n = 6/8*; PAK2 mKO, *n = 6/9*; 1/m2 dKO, *n = 6/8*. For EDL, one data point from 1/m2 dKO contraction-stimulated pTBC1D1 S237 is missing due to a lack of sample. For total protein expression, the number of determinations in each group: Control, *n = 10/10* (soleus/EDL); PAK1 KO, *n = 12/11*; PAK2 KO, *n = 13/13*; 1/m2 dKO, *n = 13/13*. Data are presented as mean ± S.E.M. with individual data points shown. A.U., arbitrary units.

## Discussion

The present study is, to our knowledge, the first to investigate the requirement of PAK1 and PAK2 in contraction-stimulated glucose transport in mouse skeletal muscle. By undertaking a systematic investigation, including pharmacological as well as genetic interventions, we show that contraction-stimulated glucose transport in isolated skeletal muscle partially requires PAK2, but not PAK1, in glycolytic EDL muscle.

In the current study, IPA-3 attenuated the increase in muscle glucose transport in response to electrically-stimulated contraction in both soleus and EDL muscle, whereas genetically targeted knockout revealed an effect of PAK2 in glycolytic EDL only. It is not unusual that pharmacological inhibition and genetically targeted mutations produce different phenotypes ^25^. This likely means that the effect of the IPA-3 on glucose transport in soleus is unspecific or alternatively, that the absent effect of genetic ablation of PAK1 and/or PAK2 in soleus is due to compensation by other mechanisms. It is important to stress that any possible compensatory mechanisms cannot be via redundancy with PAK3, as even in 1/m2 dKO muscle, PAK3 cannot be detected at the protein level ^20^.

The limited role of group I PAKs in contraction-induced glucose transport is in accordance with our recent finding that group I PAKs were largely dispensable for insulin-stimulated glucose transport in isolated mouse skeletal muscle with only a modest reduction in EDL muscles lacking PAK2 ^22^. Thus, group I PAKs are not major essential components in the regulation of muscle glucose transport. Based on recent emerging evidence, the role for group I PAKs in skeletal muscle seems instead to be related to myogenesis and muscle mass regulation ^20,23^. Additionally, in embryonic day 18.5 diaphragm, combined genetic ablation of PAK1 and PAK2 was associated with reduced acetylcholine receptor clustering at the neuromuscular junction ^20^ suggesting defects in the neuromuscular synapses.

This relatively modest requirement of group I PAKs in contraction-induced muscle glucose uptake is in contrast to the marked glucoregulatory role of Rac1 ^15,17,18^, the upstream regulator of group I PAKs. Rac1 is an essential component in the activation of the reactive oxygen-producing NADPH oxidase (NOX)-2 complex ^26,27^. Recently, it was reported that NOX2 is required for exercise-stimulated glucose uptake ^28^. Moreover, it was shown that exercise-induced NOX2 activation was completely abrogated in TA from muscle-specific Rac1 KO mice ^28^, suggesting that Rac1 mainly regulates muscle glucose uptake through activation of NOX2 in response to exercise. Alternatively, the Ral family GTPase, RalA could signal downstream of Rac1. Overexpression of a constitutively activated Rac1 mutant activated RalA in L6 myotubes ^29^ and GLUT4 translocation induced by a constitutively active Rac1 mutant was attenuated in L6-GLUT4myc myoblasts upon RalA knockdown ^29^. The RalA GTPase-activating protein GARNL1 is phosphorylated in response to in situ contraction of mouse muscle ^30^, but so far no linkage between Rac1 and RalA has been reported in relation to contraction-stimulated glucose transport.

In conclusion, contraction-stimulated glucose transport in isolated mouse skeletal muscle partially requires PAK2, but not PAK1, in glycolytic EDL muscle. Together with our previous study showing that insulin-stimulated glucose transport also partially requires PAK2, but not PAK1 ^22^, this suggests that group I PAKs play at most a minor role in the regulation skeletal muscle glucose transport.

## Materials and Methods

### Animal experiments

All animal experiments complied with the European Convention for the protection of vertebrate animals used for experimental and other scientific purposes (No. 123, Strasbourg, France, 1985; EU Directive 2010/63/EU for animal experiments) and were approved by the Danish Animal Experimental Inspectorate. All mice were maintained on a 12:12-hour light-dark cycle and housed at 22°C (with allowed fluctuation of ±2°C) with nesting material. Female C57BL/6J mice (Taconic, Denmark) were used for the inhibitor incubation study. The mice received a standard rodent chow diet (Altromin no. 1324; Brogaarden, Denmark) and water ad libitum. The mice were group-housed.

#### Double PAK1^-/-^;PAK2^fl/fl^;MyoD^iCre/+^ mice

Double knockout mice with whole-body knockout of PAK1 and conditional, muscle-specific knockout of PAK2, PAK1^-/-^;PAK2^fl/fl^;MyoD^iCre/+^ were generated as previously described ^20^. The mice were on a mixed C57BL/6/FVB background. PAK1^-/-^;PAK2^fl/fl^;MyoD^iCre/+^ were crossed with PAK1^+/-^;PAK2^fl/fl^;MyoD^+/+^ to generate littermate PAK1^-/-^;PAK2^fl/fl^;MyoD^iCre/+^ (referred to as 1/m2 dKO), PAK1^-/-^;PAK2^fl/fl^;MyoD^+/+^ (referred to as PAK1 KO), PAK1^+/-^;PAK2^fl/fl^;MyoD^iCre/+^ (referred to as PAK2 mKO), and PAK1^+/-^;PAK2^fl/fl^;MyoD^+/+^ (referred to as controls) used for experiments as previously described ^22^. At 26-35 weeks of age, female and male mice were used for the measurement of contraction-stimulated glucose transport in isolated muscle. Number of mice in each group: Control, *n = 6/4* (female/male); PAK1 KO, *n = 6/6*, PAK2 mKO, *n = 6/7*, 1/m2 dKO, *n = 6/7*. Additional mice included for measurement of muscle mass: Control, *n = 0/0* (female/male); PAK1 KO, *n = 0/1*, PAK2 mKO, *n = 3/0*, 1/m2 dKO, *n = 2/2*. Mice received standard rodent chow diet and water ad libitum. The mice were single-caged 4-7 weeks prior to the isolation of muscles. The whole-body metabolic characteristics for this cohort of mice, including insulin and glucose tolerance, have previously been described ^22^.

### Incubation of isolated muscles

Soleus and EDL muscles were dissected from anaesthetized mice (6 mg pentobarbital sodium 100 g^-1^ body weight i.p.) and suspended at resting tension (4-5 mN) in incubations chambers (Multi Myograph System, Danish Myo Technology, Denmark) in Krebs-Ringer-Henseleit buffer with 2 mM pyruvate and 8 mM mannitol at 30°C, as described previously ^31^. Additionally, the Krebs-Ringer-Henseleit buffer was supplemented with 0.1% BSA (v/v). Isolated muscles from female C57BL/6J mice were pre-incubated with 40 µM IPA-3 (Sigma-Aldrich) or a corresponding amount of DMSO (0.11%) for 45 minutes followed by 15 minutes of electrically-stimulated contractions. Isolated muscles from whole-body PAK1 KO, PAK2 mKO, 1/m2 dKO, or littermate controls were pre-incubated 10-20 minutes followed by 15 minutes of electrically-stimulated contractions. Contractions were induced by electrical stimulation every 15 sec with 2-sec trains of 0.2 msec pulses delivered at 100 Hz (∼35V) for 15 minutes. 2DG transport was measured together with 1 mM 2DG during the last 10 min of the contraction stimulation period using 0.60-0.75 µCi mL^-1^ [^3^H]-2DG and 0.180-0.225 µCi mL^-1^ [^14^C]-mannitol radioactive tracers (Perkin Elmer) as described previously ^31^. Tissue-specific [^3^H]-2DG accumulation with [^14^C]-mannitol as an extracellular marker was determined as previously described ^32^.

### Protein extraction

All muscles were homogenized 2 × 30 sec at 30 Hz using a Tissuelyser II (Qiagen, USA) in ice-cold homogenization buffer (10% (v/v) Glycerol, 1% (v/v) NP-40, 20 mM Na-pyrophosphate, 150 mM NaCl, 50 mM HEPES (pH 7.5), 20 mM β-glycerophosphate, 10 mM NaF, 2mM PMSF, 1 mM EDTA (pH 8.0), 1 mM EGTA (pH 8.0), 2 mM Na3VO4, 10 µg mL^-1^ Leupeptin, 10 µg mL^-1^ Aprotinin, 3 mM Benzamidine). After rotation end-over-end for 30 min at 4°C, lysate supernatants were collected by centrifugation (10,854-15,630 x g) for 15-20 min at 4°C.

### Immunoblotting

Lysate protein concentration was determined using the bicinchoninic acid method using bovine serum albumin (BSA) standards and bicinchoninic acid assay reagents (Pierce). Immunoblotting samples were prepared in 6X sample buffer (340 mM Tris (pH 6.8), 225 mM DTT, 11% (w/v) SDS, 20% (v/v) Glycerol, 0.05% (w/v) Bromphenol blue). Protein phosphorylation (p) and total protein expression were determined by standard immunoblotting technique loading equal amounts of protein. The polyvinylidene difluoride membrane (Immobilon Transfer Membrane; Millipore) was blocked in Tris-Buffered Saline with added Tween20 (TBST) and 2% (w/v) skim milk or 3% (w/v) BSA protein for 15 minutes at room temperature, followed by incubation overnight at 4°C with a primary antibody (Table 1). Next, the membrane was incubated with horseradish peroxidase-conjugated secondary antibody (Jackson Immuno Research) at 4°C overnight. Total ACC was detected without the use of antibodies. Instead, the membrane was incubated with horseradish peroxidase-conjugated streptavidin (P0397; Dako; 1:3000, 3% BSA) at 4°C overnight. Bands were visualized using Bio-Rad ChemiDocTM MP Imaging System and enhanced chemiluminescence (ECL+; Amersham Biosciences). Coomassie brilliant blue staining was used as a loading control ^33^. Densitometric analysis was performed using Image LabTM Software, version 4.0 (Bio-Rad, USA; RRID: SCR_014210).

**Table 1:** Antibody Table. ^#^Mouse nomenclature was used for pACC1/2 S79/212 (equivalent to human S80/221) and pTBC1D1 S231 (equivalent to human S237).

| Antibody name | Antibody ID (RRID) | Manufacturer; Catalog Number; | Species Raised in; Monoclonal or Polyclonal | Antibody dilution | Blocking buffer |
| --- | --- | --- | --- | --- | --- |
| pACC1/2 S79/212<sup>#</sup> | AB_330337 | Cell Signaling Technology; 3661 | Rabbit; Polyclonal antibody | 1:500 | 2% milk |
| Actin | AB_476693 | Sigma-Aldrich; A2066 | Rabbit; Polyclonal antibody | 1:10,000 | 2% milk |
| AMPKα2 | AB_2169716 | Santa Cruz Biotechnology; sc-19131 | Goat; Polyclonal antibody | 1:1000 | 2% milk |
| pAMPK T172 | AB_330330 | Cell Signaling Technology; 2531 | Rabbit; Polyclonal antibody | 1:1000 | 2% milk |
| GLUT4 | AB_2191454 | Thermo Fisher Scientific; PA1-1065 | Rabbit; Polyclonal antibody | 1:1000 | 2% milk |
| HKII | AB_2295219 | Santa Cruz Biotechnology; Sc-130358 | Mouse; Monoclonal antibody | 1:1000 | 2% milk |
| PAK1 | AB_330222 | Cell Signaling Technology; 2602 | Rabbit; Polyclonal antibody | 1:500 | 2% milk |
| PAK2 | AB_2283388 | Cell Signaling Technology; 2608 | Rabbit; Polyclonal antibody | 1:500 | 2% milk |
| TBC1D1 | AB_2814949 | Abcam; ab229504 | Rabbit; Polyclonal | 1 µg/µL | 2% milk |
| pTBC1D1 S231<sup>#</sup> | AB_10807809 | Millipore; 07-2268 | Rabbit; Polyclonal antibody | 1:1000 | 2% milk |

### Statistical analyses

Data are presented as mean ± S.E.M. or when applicable, mean ± S.E.M. with individual data points shown. Statistical tests varied according to the dataset being analyzed and the specific tests used are indicated in the figure legends. Datasets were normalized by square root, log10 or inverse transformation if not normally distributed or failed equal variance test. If the null hypothesis was rejected, Tukey’s post hoc test was used to evaluate significant main effects of genotype and significant interactions in ANOVAs. P < 0.05 was considered statistically significant. P<0.1 was considered a tendency. Except for mixed-effects model analyses performed in GraphPad Prism, version 8.2.1. (GraphPad Software, La Jolla, CA, USA; RRID: SCR_002798), all statistical analyses were performed using Sigma Plot, version 13 (Systat Software Inc., Chicago, IL, USA; RRID: SCR_003210). Due to missing data points, differences between genotypes and the effect of electrically-stimulated contraction were assessed with a mixed-effects model analysis in Fig. 2H+I.

## Data availability

The datasets generated and analyzed during the current study are available from the corresponding author upon reasonable request. No novel applicable resources were generated or analyzed during the current study.

## Acknowledgements

We thank our colleagues at the Section of Molecular Physiology, Department of Nutrition, Exercise, and Sports (NEXS), Faculty of Science, University of Copenhagen, for fruitful discussions on this topic. We acknowledge the skilled technical assistance of Betina Bolmgren and Mona Ali (Section of Molecular Physiology, NEXS, Faculty of Science, University of Copenhagen, Denmark). The PAK1/m2 dKO founder mice were a kind gift from Giselle A. Joseph and Robert S. Krauss (Department of Cell, Developmental, and Regenerative Biology, Icahn School of Medicine at Mount Sinai, New York, USA).

## Author contributions

**LLVM:** Conceptualization; Methodology; Formal analysis; Investigation; Writing - Original Draft; Writing - Review & Editing; Visualization; Project administration; Funding acquisition. **ILN:** Investigation; Writing - Review & Editing. **JRK:** Investigation; Writing - Review & Editing; Funding acquisition. **NRA:** Investigation; Writing - Review & Editing. **TEJ:** Investigation; Writing - Review & Editing; Funding acquisition. **LS:** Conceptualization; Methodology; Investigation; Writing - Original Draft; Writing - Review & Editing; Supervision; Project administration; Funding acquisition. **EAR:** Conceptualization; Methodology; Writing - Original Draft; Writing - Review & Editing; Supervision; Project administration; Funding acquisition. EAR is the guarantor of this work and takes responsibility for the integrity of the data and the accuracy of the data analysis.

